# Plasma Gelsolin Prevents Organ-Specific Inflammation and Enhances Innate Immune Function in a Systemic *Candida albicans* Infection

**DOI:** 10.1101/2025.09.22.677758

**Authors:** Łukasz Suprewicz, Alicja Walewska, Andrzej Namiot, Sylwia Deptuła-Chmielewska, Paul A. Janmey, Robert Bucki

## Abstract

Systemic *Candida albicans* infections (candidemia) remain a significant cause of morbidity and mortality in immunocompromised patients, mainly due to severe inflammation, organ damage, and delayed fungal clearance. Plasma gelsolin (pGSN), an actin-binding protein with immunomodulatory properties, has demonstrated protective effects in bacterial sepsis models, but its role in fungal infections remains unexplored. Here, we evaluated the impact of pGSN in a murine model of candidemia. Mice intravenously challenged with *C. albicans* exhibited strong inflammatory responses, particularly in the kidneys and lungs, as visualized by IRDye 800CW 2-deoxyglucose imaging and confirmed by histopathological examination. Subcutaneous injection of pGSN significantly reduced systemic and organ-specific inflammation, decreased blood fungal burden, and prevented microabscess formation in the kidneys. In parallel, pGSN suppressed inflammatory gene expression in whole blood and enhanced phagocytic activity of human monocytes. Additionally, pGSN modulated monocyte reactive radical production by increasing nitric oxide and reducing hydrogen sulfide and reactive oxygen species production. These results highlight the dual immunomodulatory and host-protective properties of pGSN, supporting its potential as an adjunctive therapy in fungal sepsis.

## 1. INTRODUCTION

Systemic candidiasis, most commonly caused by *Candida albicans* (*C. albicans*), remains a major cause of morbidity and mortality, particularly in immunocompromised patients and those receiving broad-spectrum antibiotics or invasive medical interventions (1, 2). Despite the availability of antifungal therapies, mortality rates for candidemia can exceed 40%, especially in patients with delayed diagnosis or organ failure (3). The pathogenesis of systemic *Candida* infection is not solely attributed to fungal proliferation, but also to the host’s dysregulated immune response, which drives tissue damage and multiorgan dysfunction (4).

One of the major challenges in treating candidemia is the dual requirement for fungal clearance and control of inflammation. While antifungal drugs can suppress fungal growth, they do not address the collateral inflammatory damage induced by excessive cytokine release, immune cell infiltration, and microvascular dysfunction (5). Moreover, host immune responses vary across organs, with the kidneys exhibiting the most severe inflammation and tissue injury during fungal sepsis (6–8). Therefore, there is a critical need for adjunctive therapies that can modulate the host immune response without compromising pathogen clearance.

Plasma gelsolin (pGSN) is a circulating actin-binding protein that plays a crucial role in maintaining immune homeostasis, scavenging extracellular actin, modulating Toll-like and scavenger receptor signaling, and preserving endothelial integrity (9–13). Under conditions of acute inflammation, including sepsis and trauma, pGSN levels drop significantly, impairing its protective functions (14). Restoring pGSN levels has been shown to mitigate inflammatory damage and improve survival in preclinical models of bacterial sepsis and acute lung injury (15, 16). In murine models of bacterial infection, pGSN supplementation has been shown to reduce cytokine storm, enhance bacterial clearance, and improve survival outcomes (17, 18). Furthermore, early-phase clinical studies have reported that exogenous pGSN administration is safe and may be beneficial in critically ill patients with pneumonia or sepsis (19–21). However, its role in fungal infections, particularly in the context of systemic *C. albicans* infection, remains unexplored.

In this study, we investigated the effects of recombinant human pGSN in a murine model of systemic candidiasis. We hypothesized that subcutaneous administration of pGSN would attenuate inflammation, reduce fungal burden, and support innate immune function. Using a combination of *in vivo* imaging, histopathology, gene expression profiling, and human monocyte assays, we provide evidence that pGSN acts as a dual-function host-directed therapy by modulating both inflammation and antifungal immunity.

## 2. MATERIALS AND METHODS

### 2.1. Plasma gelsolin

The recombinant human plasma gelsolin used in our study was expressed in *E. coli* cells and provided by BioAegis Therapeutics (North Brunswick, USA).

### 2.2. Inoculum preparation

The *C. albicans* reference strain (1408) was obtained from the Polish Collection of Microorganisms (Polish Academy of Sciences, Wroclaw, Poland) and stored in glycerol stocks at −80°C. For experimental use, the strain was plated on Sabouraud dextrose agar with chloramphenicol (Lab-Agar; Biomaxima, Lublin, Poland) and grown at 37°C. Yeast cells were counted using a hemocytometer and diluted to the desired concentration in sterile 0.9% NaCl for *in vivo* use or in cell culture medium for *in vitro* experiments.

### 2.3. Murine model of candidemia

Animal procedures were approved by the Local Ethics Committee for Animal Experimentation in Olsztyn, Poland (approval numbers 66/2020 and 12/2024). Female CAnN.Cg-Foxn1/CrL mice (10 weeks old; Charles River) were randomized into four experimental groups (**Table 1**). All mice were screened for bacterial, viral, and parasitic infections prior to the study (Alab Bioscience, Poland). Mice received an intravenous tail vein injection of *C. albicans* (10 CFU/mL) or 0.9% NaCl, each combined with IRDye 800CW 2-deoxyglucose (10 nM, #926-08946, Li-Cor). At 12 hours post-injection, animals were administered subcutaneously with either pGSN (10 mg/mL, 100 μL) or 0.9% NaCl. Whole-body near-infrared imaging was performed at 12, 16, 20, and 24 hours post-infection. Core body temperature measurements were taken using a handheld infrared scanner every 4 hours from the start of the experiment. At 24 hours, mice were sacrificed, and blood and organs were collected. Blood was drawn via cardiac puncture into EDTA tubes for analysis of fungal burden and gene expression. Organs were fixed in 10% formaldehyde and left for imaging and hematoxylin and eosin (H&E) staining for histopathological evaluation. All solutions were sterile-filtered using 0.2 μm syringe filters.

### 2.4. Imaging of inflammatory response

IRDye 800CW 2-deoxyglucose was used to assess inflammation. At the indicated time points, animals were anesthetized with isoflurane and scanned using the Pearl Trilogy Imaging System (Li-Cor). After sacrifice, formalin-fixed organs were imaged *ex vivo* using the Odyssey CLx Imager (Li-Cor) to assess organ-specific fluorescence intensity.

### 2.5. Fungal outgrowth in the blood

Blood samples were serially diluted and plated on Sabouraud agar for quantification of fungal burden. Plates were incubated at 38°C for 48 hours before colony counts were performed.

### 2.6. RT-qPCR

Total RNA was extracted from 100 μL of blood using the Universal RNA Purification Kit, according to the manufacturer’s instructions (E3598-02; EURx, Gdansk, Poland). RNA concentration and purity were assessed using a Qubit 4 fluorometer (Thermo Fisher Scientific). cDNA synthesis was performed with the iScript cDNA Synthesis Kit (1708891; Bio-Rad). qRT-PCR was carried out using 5 ng of cDNA and SsoAdvanced Universal SYBR Green Supermix (1725274; Bio-Rad) in a 20 μL reaction. Primers used are listed in **Table S1**. Reactions were run on the CFX Opus 96 Real-Time PCR Detection System (Bio-Rad) using the following program: 95°C for 2 min, followed by 40 cycles of 95°C for 5 s and 60°C for 30 s. GAPDH served as the reference gene. Relative gene expression was calculated using the 2^-ΔΔCt^ method.

### 2.7. Human monocyte isolation

Peripheral blood was collected from healthy donors under the approval of the Bioethics Committee at the Medical University of Bialystok (APK.002.234.2021). Monocytes were isolated using the Dynabeads CD14+ positive isolation kit (11367D; Invitrogen) according to the manufacturer’s protocol. Cells were counted using a hemocytometer and resuspended in serum- and antimycotic-free RPMI 1640 medium (ATCC) to avoid interference from exogenous pGSN. Cells were maintained at 37°C with 5% CO_2_.

### 2.8. Phagocytosis assay

Monocytes (1 × 10 cells/well) were seeded in 96-well plates pre-coated with 0.01% poly-L-lysine (Sigma-Aldrich). After monocyte serum starvation for 2 hours, calcofluor white-stained *C. albicans* (0.25 μg/mL for 30 min, RT) was added at a multiplicity of infection (MOI – a number of fungal cells per monocyte) of 5. After 2 hours of incubation at 37°C, the cells were washed with PBS, fixed in 3.7% paraformaldehyde for 15 minutes, permeabilized with 0.1% Triton X-100 for 10 minutes, and blocked with 0.1% BSA for 30 minutes. Cells were stained with Alexa Fluor 647-conjugated wheat germ agglutinin (WGA; 5 μg/mL; W32466, Invitrogen) for 10 minutes, mounted in antifade medium (Abcam), and imaged using confocal microscopy (Leica DMI8). Phagocytic activity was assessed as (i) percentage of engaged cells (monocytes adhering to or internalizing fungi) and (ii) phagocytic index (fungal cells internalized per 100 monocytes). At least eight randomly selected fields were analyzed per condition across four independent experiments.

### 2.9. Reactive species assessment

Monocytes (2 × 10 cells/well) were plated in 96-well plates and serum-starved for 2 hours. Cells were treated with *C. albicans* (MOI 5), pGSN, and their combination for 2 hours, followed by incubation with redox-sensitive probes: DAF-2 (5 nM; nitric oxide), P3 (50 nM; hydrogen sulfide), and DCFH-DA (20 μM; reactive oxygen species) (Sigma-Aldrich). Fluorescence was recorded at 488/535 nm using a Varioskan Lux microplate reader (Thermo Fisher Scientific). Background fluorescence was subtracted, and data were normalized to untreated controls.

### 2.10. Statistics

Statistical analyses were performed using OriginPro 2021 (OriginLab Corporation, Northampton, MA, USA). The number of replicates, statistical test used, and significance levels are indicated in the figure legends. Data are presented as mean ± standard deviation (SD), unless otherwise stated.

## 3. RESULTS

### 3.1. pGSN reduces systemic inflammation and circulating fungal burden in a murine model of candidemia

To investigate the therapeutic potential of pGSN in systemic *C. albicans* infection, we established a murine model of candidemia using athymic nude mice (CAnN.Cg-Foxn1 <NU>/CrL), which lack functional T cells and therefore allow us to study mostly innate immunity, which is often imbalanced in immunocompromised patients at highest risk for invasive fungal disease. Mice were randomized into four experimental groups receiving either *C. albicans* or NaCl intravenously, with or without subcutaneous pGSN treatment 12 hours post-infection (**Table 1 and Fig. 1A**).

**Table 1.**
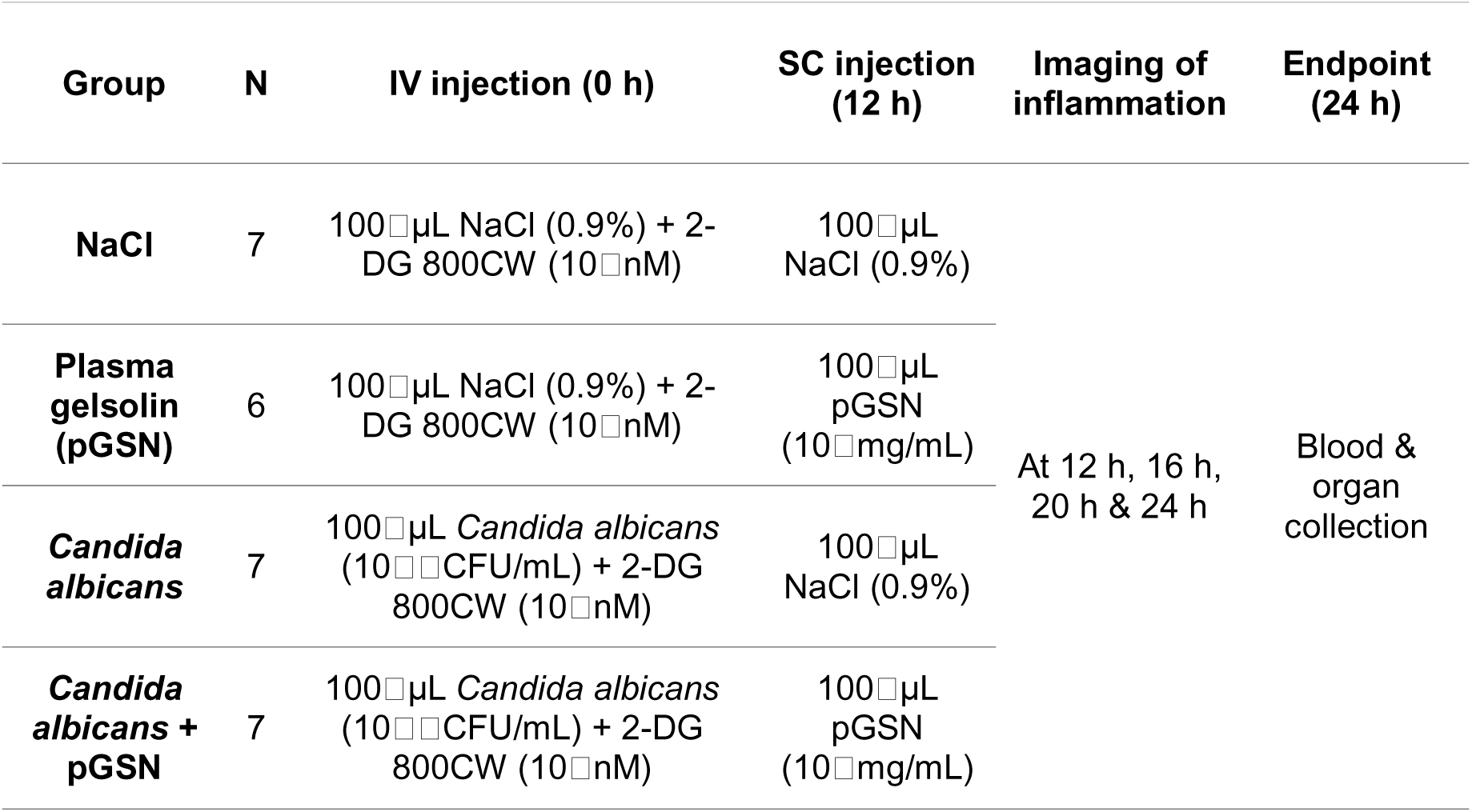
Experimental setup for the systemic *Candida albicans* infection model with plasma gelsolin intervention.

**Figure 1.**
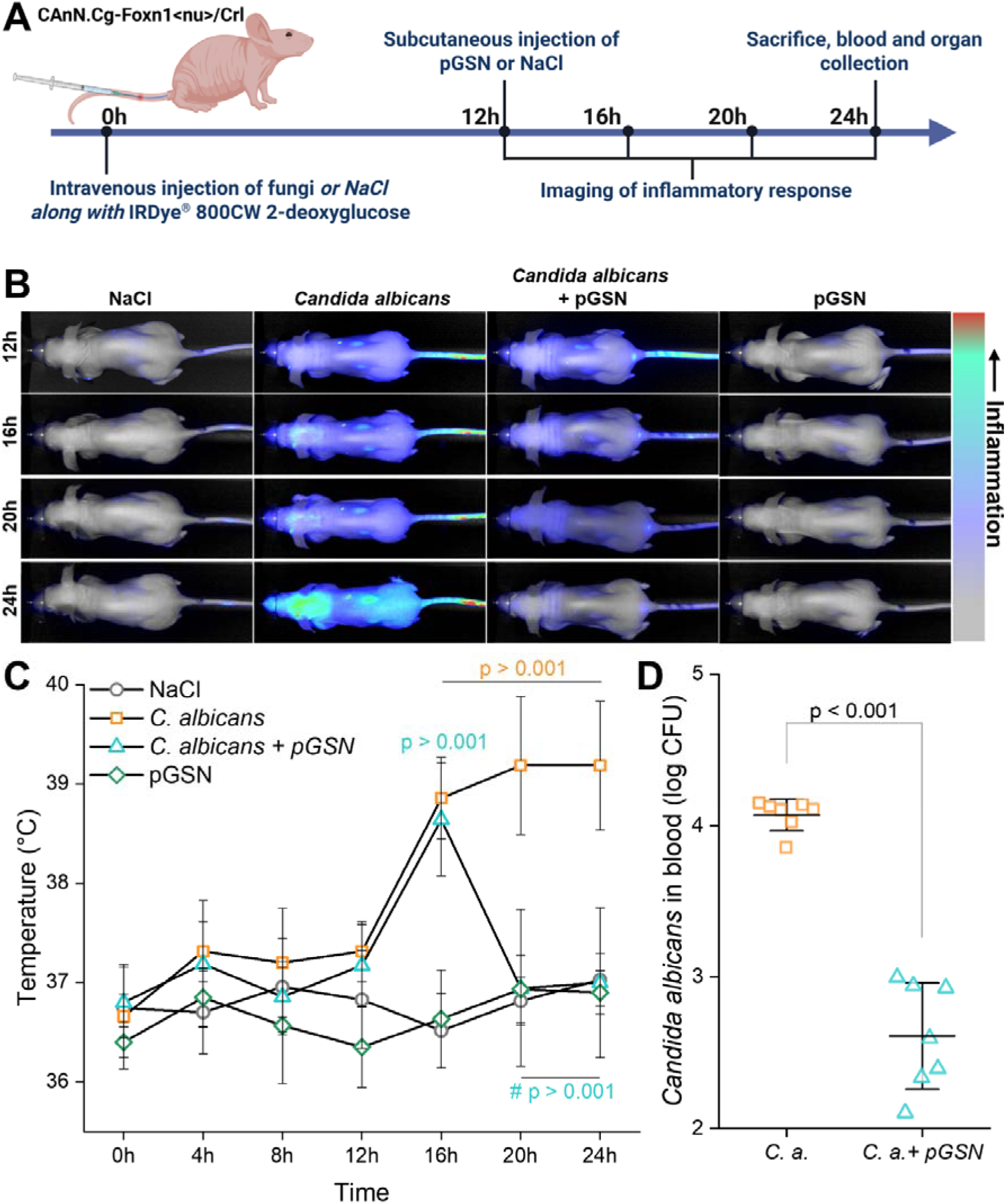
Plasma gelsolin diminished the inflammatory response caused in a murine model of candidemia. (**A**) Schematic overview of the experimental workflow. Female CAnN.Cg-Foxn1 <NU>/CrL mice (10 weeks old) were intravenously injected with *Candida albicans* (10 CFU/mL) or NaCl, along with IRDye 800CW 2-deoxyglucose (10 nM). At 12 hours post-injection, animals received a subcutaneous dose of either pGSN (10 mg/mL) or NaCl. Mice were imaged at 12, 16, 20, and 24 hours and sacrificed at 24 hours for sample collection. (**B**) Representative whole-body near-infrared images of mice from each treatment group acquired at 12, 16, 20, and 24 hours post-injection of fungal cells. The color scale indicates the relative intensity of the inflammatory signal. (**C**) Core body temperature of mice over the 24-hour course, showing elevated temperature in Candida albicans-infected animals and mitigation in the pGSN-treated group. (**D**) Fungal burden (log CFU) in blood at 24 hours post-infection. Data are presented as mean±SD. Statistical significance was determined by one-way ANOVA with post-hoc Tukey’s test (**C**) or an unpaired Student’s t-test (**D**). n = 6–7 per group.

To monitor systemic inflammation *in vivo*, we co-injected mice with 800CW 2-deoxyglucose (2DG-800CW), a near-infrared glucose analog that accumulates in metabolically active, inflamed tissues (15). Serial imaging over a 12-hour period revealed a progressive increase in whole-body fluorescence in *C. albicans*-infected animals, with a notable signal localized to the abdominal region, corresponding to the kidneys. In contrast, pGSN-treated animals displayed a visibly reduced inflammatory signal throughout the imaging time course (**Fig. 1B**).

In parallel, we measured core body temperature to assess systemic inflammation. *C. albicans* infection induced a significant rise in temperature compared to NaCl controls, whereas pGSN treatment decreased this response, maintaining values closer to baseline (**Fig. 1C**).

To determine the effect of pGSN on fungal dissemination, we quantified fungal burden in the blood 24 hours after infection. Mice receiving pGSN exhibited a marked reduction in colony-forming units (CFU) in blood compared to untreated *C. albicans*-infected animals (**Fig. 1D**), indicating pGSN’s ability to reduce systemic inflammation and increase fungal clearance.

### 3.2. pGSN protects against organ-specific inflammation during systemic candidiasis

To assess the distribution and severity of inflammation at the organ level, we performed *ex vivo* imaging of dissected organs 24 hours post-infection. In *C. albicans*-infected animals, the strongest fluorescence signal was observed in the liver, lungs and kidneys (**Figure 2A**). Quantitative analysis of fluorescence intensities across all individual animals confirmed these trends (**Fig. 2B and S2**). pGSN-treated *C. albicans*-infected mice showed significantly lower inflammatory activity in lungs and kidneys compared to untreated infected mice. At the same time, smaller changes were observed in the liver, spleen, or heart. As expected, a high baseline 2DG signal was also detected in the liver and kidneys across all groups, including the NaCl and pGSN controls, due to the physiological metabolism of 2DG, as well as the clearance and excretion of the fluorescent probe (22–24). This background signal was consistent among animals (**Fig. S1**). These findings indicate that pGSN diminished inflammation at key target sites of fungal dissemination.

**Figure 2.**
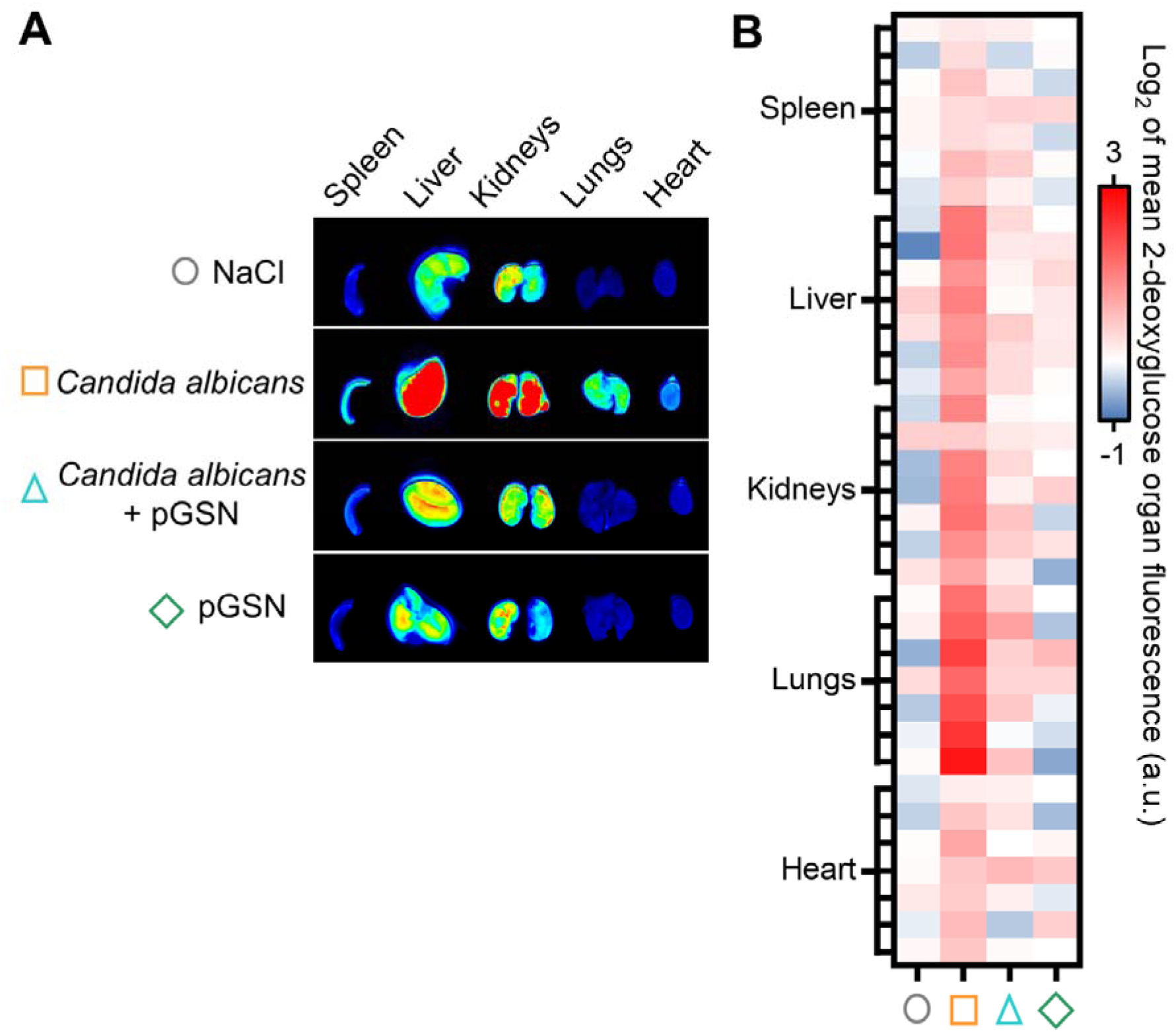
Plasma gelsolin reduces organ-specific inflammatory burden in systemic candidiasis. (**A**) *Ex vivo* near-infrared fluorescence imaging of isolated organs (spleen, liver, kidneys, lungs, and heart) collected at 24 hours post-intravenous injection of *Candida albicans* or NaCl. Mice received either plasma gelsolin (pGSN) or NaCl subcutaneously at 12 hours. Representative fluorescence images show elevated glucose uptake in the liver, kidneys, and lungs of infected animals, indicating tissue inflammation. (**B**) Heatmap showing log_2_-transformed mean fluorescence intensities of 2-deoxyglucose in each organ across all individual animals. Each column represents a single animal; n = 6–7 per group.

To evaluate the extent of organ damage during candidemia, formalin-fixed lungs, kidneys, and liver were processed for hematoxylin and eosin (H&E) staining and assessed by a pathologist. Consistent with the *ex vivo* imaging data, *C. albicans* infection induced strong tissue-specific inflammation in the kidneys and lungs, while the liver remained largely unaffected (**Table 2**).

**Table 2.**
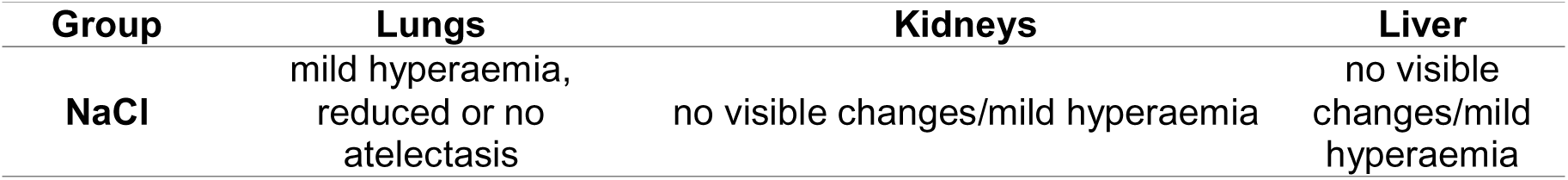
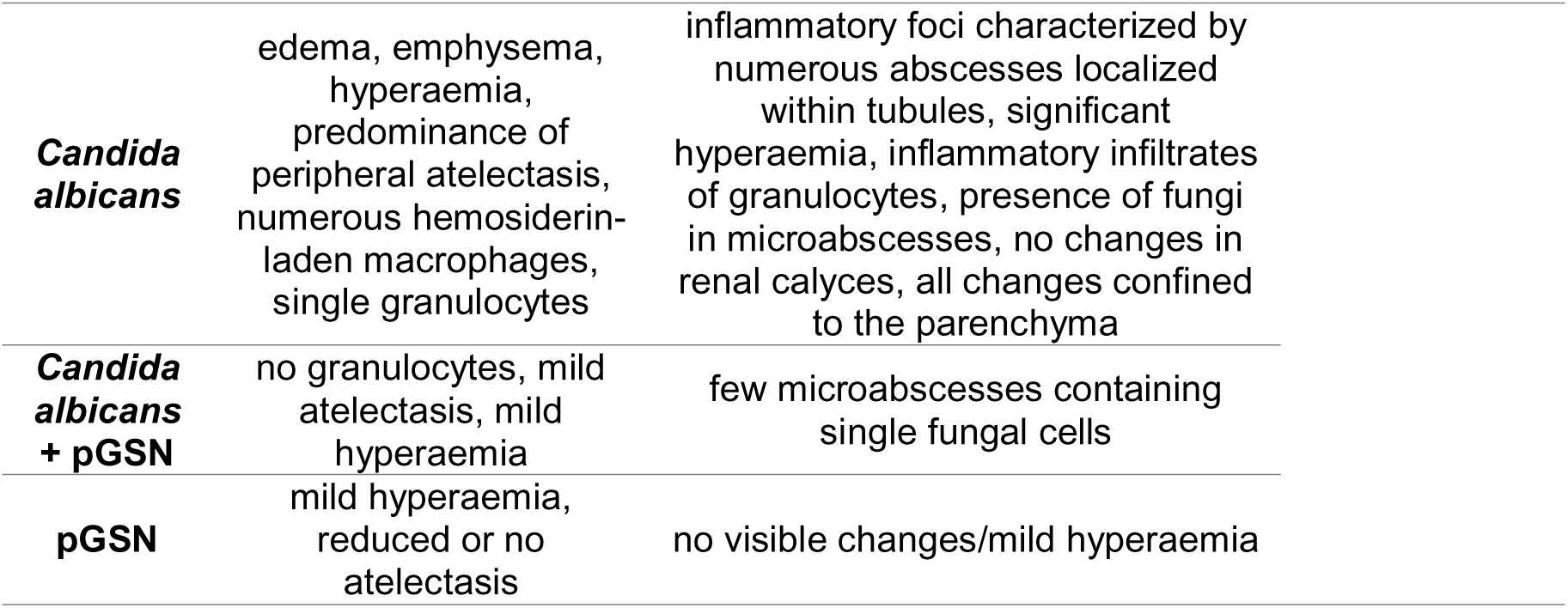
Pathological changes in major organs 24 hours after the initiation of treatment.

| Group | Lungs | Kidneys | Liver |
| --- | --- | --- | --- |
| NaCl | mild hyperaemia,<br>reduced or no<br>atelectasis | no visible changes/mild hyperaemia | no visible<br>changes/mild<br>hyperaemia |

|  |  |  |
| --- | --- | --- |
| ***Candida albicans*** | edema, emphysema, hyperaemia, predominance of peripheral atelectasis, numerous hemosiderin-laden macrophages, single granulocytes | inflammatory foci characterized by numerous abscesses localized within tubules, significant hyperaemia, inflammatory infiltrates of granulocytes, presence of fungi in microabscesses, no changes in renal calyces, all changes confined to the parenchyma |
| ***Candida albicans* + pGSN** | no granulocytes, mild atelectasis, mild hyperaemia | few microabscesses containing single fungal cells |
| **pGSN** | mild hyperaemia, reduced or no atelectasis | no visible changes/mild hyperaemia |

Kidney sections from infected mice revealed extensive inflammatory foci characterized by tubular microabscesses, marked granulocytic infiltration, and the presence of fungal cells within abscesses (**Figs. 3 and 4**). These lesions were primarily confined to the renal parenchyma, with no apparent involvement of the renal calyces. In contrast, mice treated with pGSN exhibited only sparse microabscesses, rarely containing single fungal cells, and lesser granulocytic infiltration.

**Figure 3.**
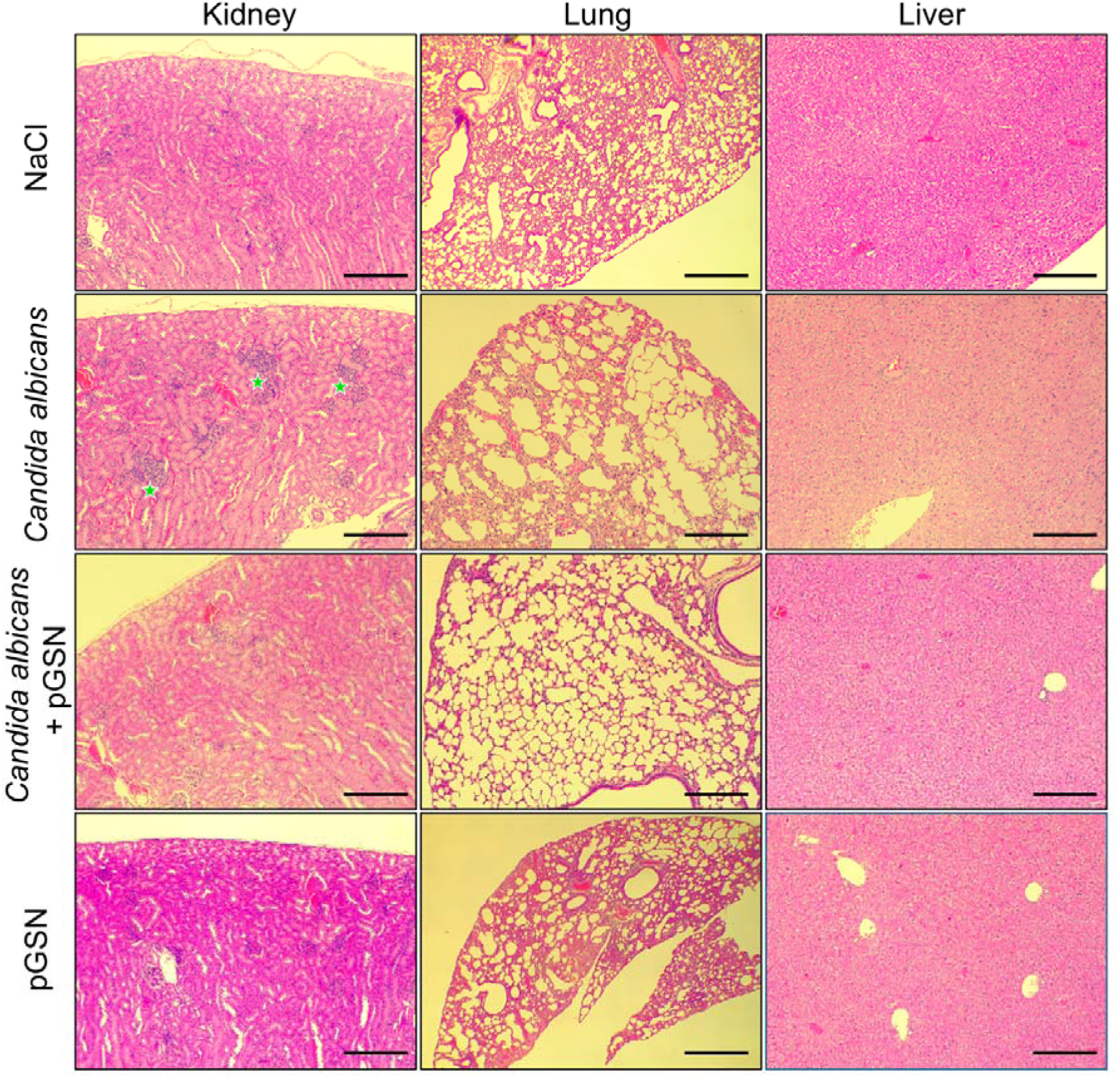
Histopathological analysis of kidney, lung, and liver tissue reveals tissue-specific inflammation and the protective effect of plasma gelsolin. Representative hematoxylin and eosin (H&E) stained sections of kidney, lung, and liver harvested at 24 hours post-treatment from all experimental groups. In *Candida albicans*-infected animals, kidneys show prominent inflammatory foci with numerous microabscesses (indicated by green stars), while lungs exhibit emphysematous changes and thickened alveolar walls. No major pathology is observed in the liver. Co-administration of plasma gelsolin (pGSN) with *Candida albicans* reduces inflammation and largely prevents abscess formation. Scale bars = 200 μm.

**Figure 4.**
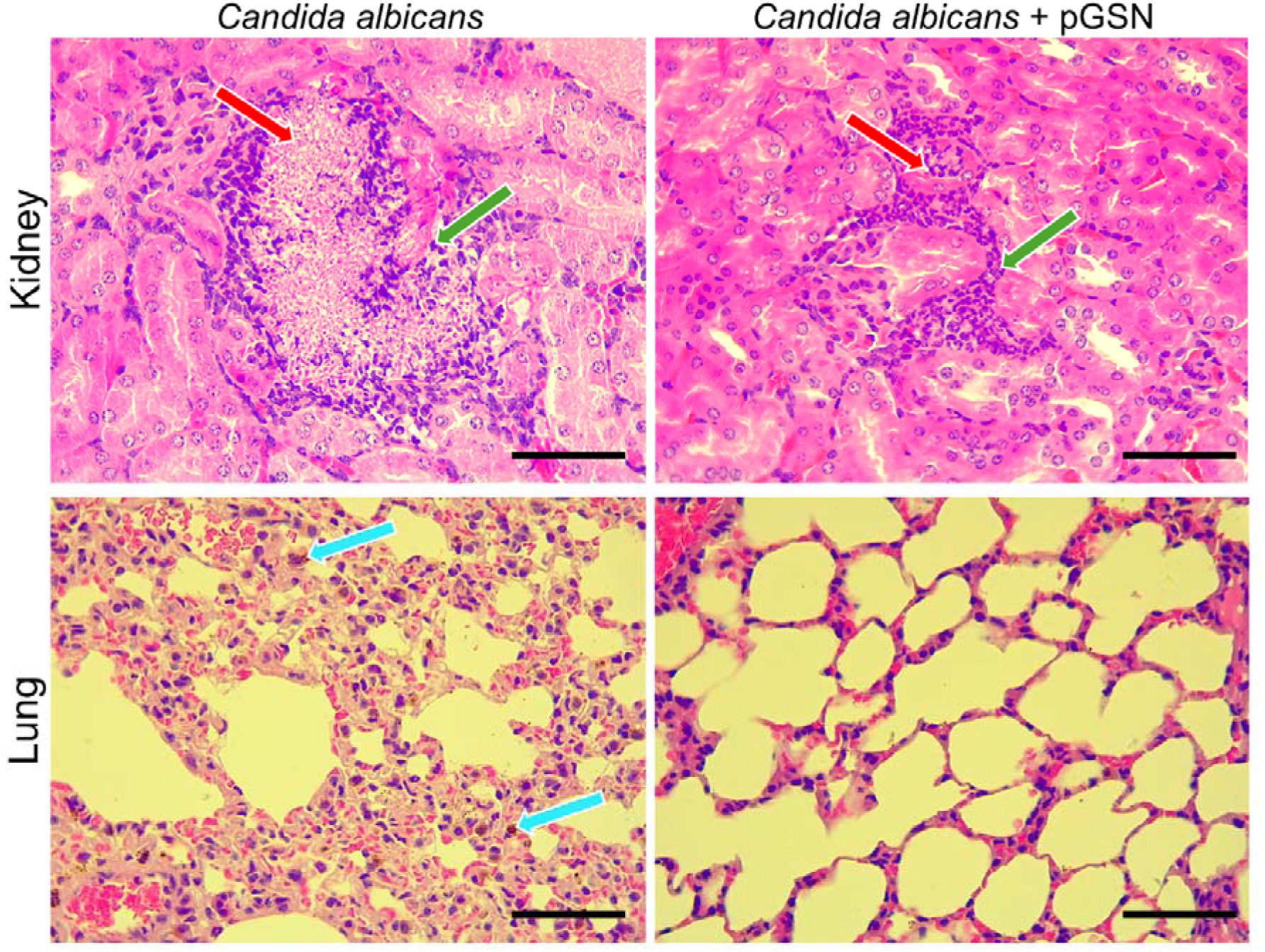
Plasma gelsolin reduces microabscess formation in kidneys and alleviates lung pathology during systemic *Candida albicans* infection. High-magnification hematoxylin and eosin (H&E) stained images of kidney and lung tissue collected at 24 hours post-intravenous injection of Candida albicans, with or without plasma gelsolin (pGSN) treatment. In the kidneys of infected animals, multiple tubular microabscesses containing fungal cells (red arrows) are characterized by granulocytic infiltration (green arrows). Co-administration of pGSN markedly reduces the number and extent of abscesses. (**B**) Lung sections from infected mice reveal alveolar collapse, hyperemia, and leukocyte infiltration. Blue arrows indicate hemosiderin-rich macrophages. These inflammatory changes are mitigated in animals treated with pGSN. Scale bars = 50 μm.

In the lungs, *Candida*-infected animals showed signs of pulmonary injury, including edema, emphysema, peripheral atelectasis, and the presence of hemosiderin-laden macrophages, indicative of tissue damage and hemorrhage (**Figs. 3 and 4**). These pathological changes were notably reduced in animals receiving pGSN, which exhibited only mild hyperemia and minimal structural disruption.

Liver tissue from all groups showed minimal to no pathological alterations. No histological abnormalities were observed in either the NaCl or pGSN-only control groups, indicating that neither vehicle nor recombinant pGSN induces baseline inflammation or damage.

Together, these histological findings confirm that pGSN mitigates *Candida*-induced inflammation and tissue injury, particularly in the kidneys, which are the primary target organs during systemic fungal infection. It can also be concluded that the gelsolin administered during the experiment does not show toxicity to hepatocytes.

### 3.3. pGSN reduces systemic inflammatory gene expression during candidemia

To assess the systemic immune response to *C. albicans* infection and the effect of pGSN treatment, we analyzed the expression of inflammatory cytokine genes in whole blood collected 24 hours post-infection. Infected animals showed a robust upregulation of several pro-inflammatory genes, including IL-1β, IL-6, TNF-α, and TGF-β (**Fig. 5A–D**). Treatment with pGSN significantly suppressed the expression of these cytokines, suggesting a systemic anti-inflammatory effect. These observations are consistent with the attenuation of both body temperature and organ-specific inflammation shown in **Figs 1 and 3**.

**Figure 5.**
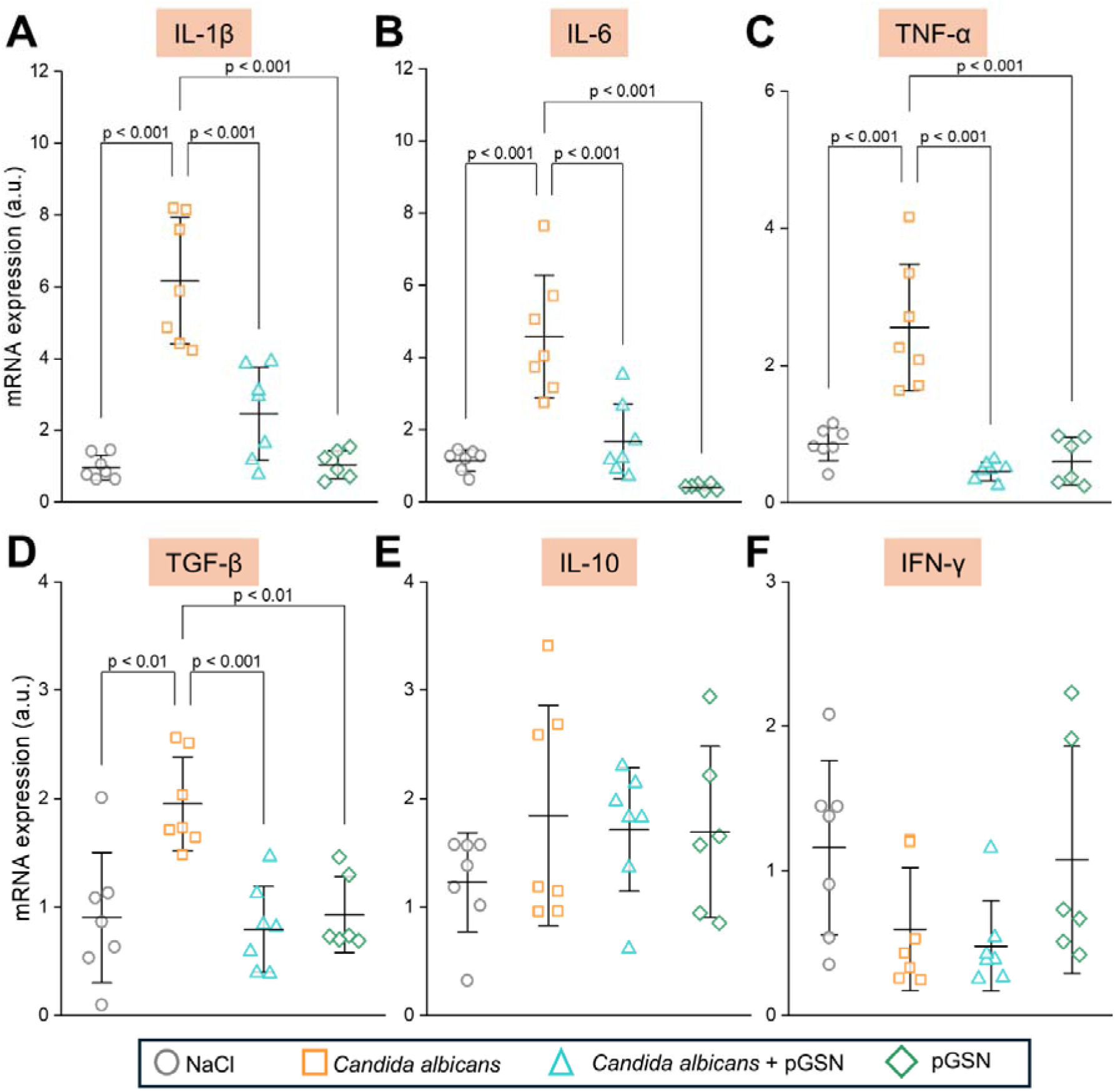
Plasma gelsolin attenuates systemic inflammatory gene expression in whole blood during candidemia. pGSN treatment reduced *Candida albicans*-driven elevated expression of several inflammatory mediators, including (**A**) IL-1β, (**B**) IL-6, (**C**) TNF-α, and (**D**) TGF-β. No statistically significant change was recorded for (**E**) IL-10 and (**F**) IFN-γ. Expression levels were normalized to housekeeping genes and are shown relative to the NaCl-only group. Data are presented as mean ± SD. Each dot represents one animal (n = 6-7). Statistical significance was determined by one-way ANOVA with post-hoc Tukey’s test.

Expression levels of IL-10, an anti-inflammatory cytokine, exhibited a non-significant upward trend in pGSN-treated animals compared to infected controls, indicating a possible shift toward resolution-phase signaling (**Fig. 5E**). Notably, downregulation in IFN-γ levels was detected across the *Candida*-infected groups (**Fig. 5F**).

These data indicate that pGSN modulates systemic inflammation at the transcriptional level during candidemia, primarily by suppressing pro-inflammatory gene expression without impairing anti-inflammatory responses.

### 3.4. pGSN enhances the antifungal activity of human monocytes

To investigate the immunomodulatory effects of pGSN on innate immune function in a human context, we examined the interaction between *C. albicans* and primary human monocytes. Peripheral blood monocytes were selected due to their abundance and clinical relevance during infections. While neutrophils have been extensively studied in antifungal defense, the role of monocytes in this context, particularly in response to pGSN, has not been well characterized (12).

Using fluorescence and confocal microscopy, we analyzed the uptake of *C. albicans* by monocytes in the presence or absence of pGSN. Cells were stained with wheat-germ agglutinin (WGA) to define monocyte membranes, while calcofluor white distinguished fungal cells. High-resolution 3D-stack imaging revealed that pGSN-treated monocytes internalize fungal cells, as visualized by the presence of engulfed *Candida* within the monocyte cytoplasm (**Fig. 6A and 6Ai).**

**Figure 6.**
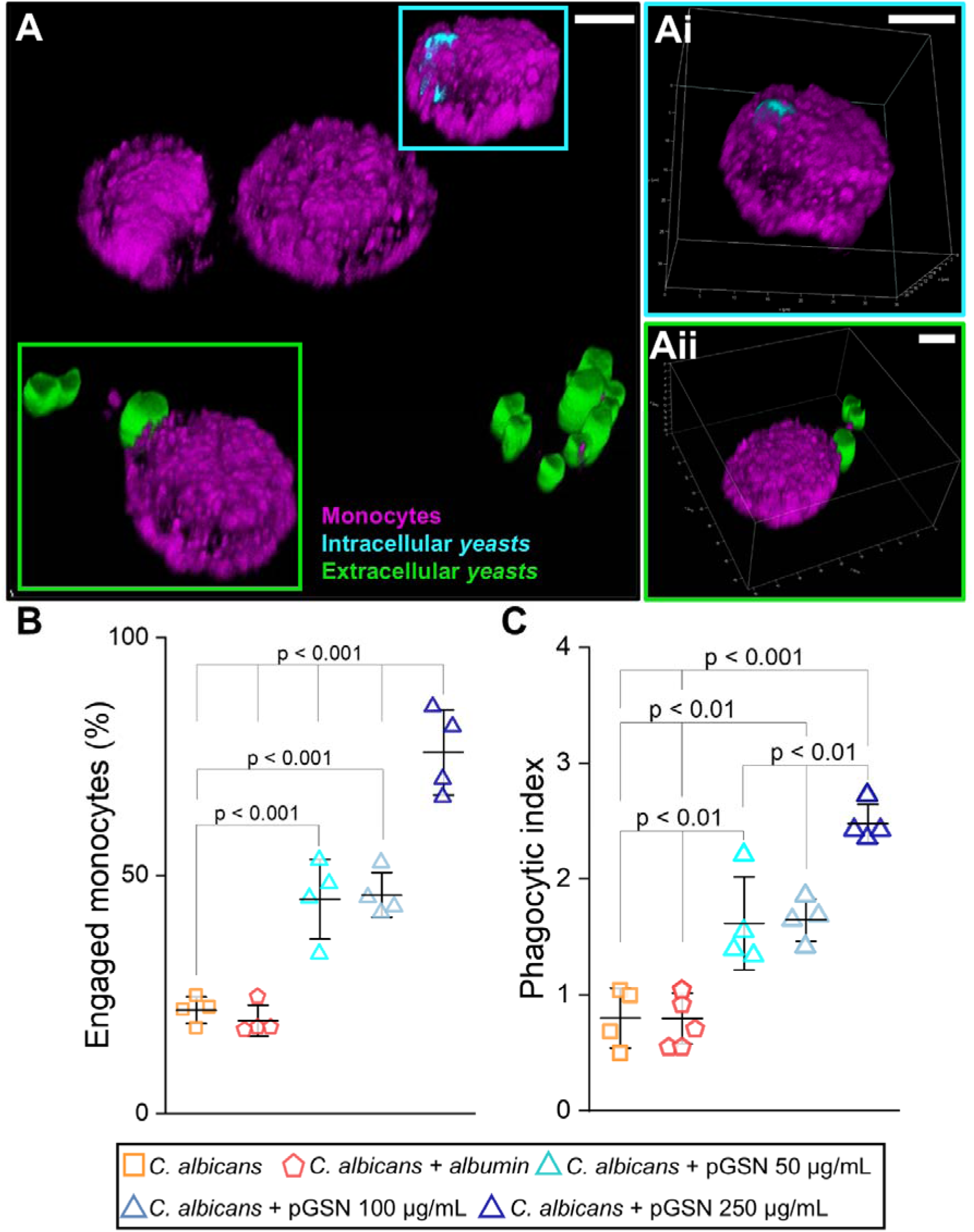
Plasma gelsolin enhances the phagocytic activity of human monocytes against *Candida albicans*. (**A**) Representative fluorescence microscopy images of monocytes (magenta) isolated from healthy donors, exposed to *Candida albicans* in the presence or absence of plasma gelsolin (pGSN). Internalized fungal cells (cyan) are visible within the monocyte cytoplasm, while extracellular yeasts are green. (**Ai**) 3D-confocal stack image showing a monocyte with an internalized Candida cell within the cytoplasm. (**Aii**) Confocal stack showing a monocyte attached to the extracellular fungal cell. (**B**) Quantification of the percentage of monocytes actively engaged in phagocytosis. (**C**) Phagocytic index calculated as the average number of *Candida albicans* cells internalized per monocyte. Data represent mean ± SD from n = 4 independent experiments. Statistical comparisons were made using one-way ANOVA with post-hoc Tukey’s test.

Quantification of phagocytic engagement showed a significant increase in the percentage of monocytes participating in phagocytosis following pGSN treatment (**Fig. 6Aii and 6B**). Additionally, the phagocytic index (the average number of fungal cells internalized per monocyte) was also elevated in the pGSN-treated condition (**Fig. 6C**), indicating enhanced uptake efficiency.

These findings suggest that pGSN not only mitigates systemic inflammation but also augments the functional antifungal capacity of monocytes, potentially contributing to the reduced fungal burden observed *in vivo*.

### 3.5. pGSN modulates reactive radical production in human monocytes during fungal challenge

Reactive radical signaling plays a critical role in regulating phagocyte function, including microbial killing, cytokine release, and resolution of inflammation (25). To assess how pGSN influences the redox state of human monocytes in the context of *C. albicans* infection, we measured the production of three key mediators: nitric oxide (NO), hydrogen sulfide (H_2_S), and reactive oxygen species (ROS).

Upon exposure to *C. albicans*, monocytes exhibited elevated production of NO, H_2_S, and ROS, reflecting their activation in response to fungal stimulation (**Fig. 7A–C**). Interestingly, pGSN treatment further increased NO levels beyond those seen with *Candida* stimulation alone, suggesting that pGSN may enhance signaling or antimicrobial activity through the NO pathway. In contrast, pGSN significantly attenuated *Candida*-induced H_2_S and ROS production, indicating selective suppression of oxidative and sulfide stress.

**Figure 7.**
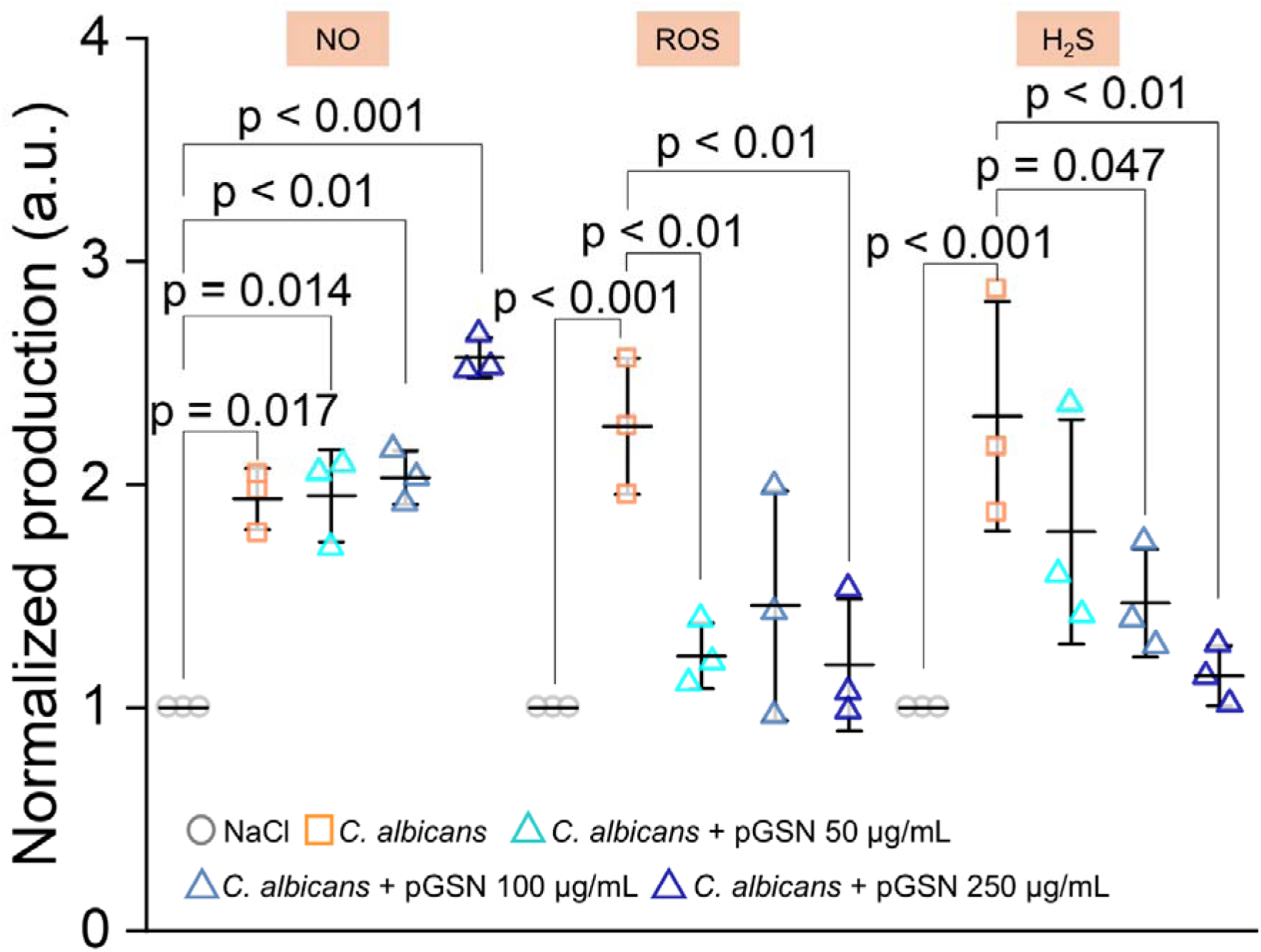
Plasma gelsolin modulates monocyte reactive radical production in response to fungal challenge. Monocytes isolated from healthy human donors were incubated with Candida albicans in the presence or absence of plasma gelsolin (pGSN). Production of (**A**) nitric oxide, (**B**) hydrogen sulfide, and (**C**) reactive oxygen species was measured using fluorescent probes. Data are presented as mean ± SD from n = 3 independent experiments. Statistical significance was determined using one-way ANOVA with post-hoc Tukey’s test.

Notably, pGSN alone increased NO levels in resting monocytes but did not significantly alter H_2_S or ROS production (**Fig. S3**). This selective modulation of reactive radical species suggests that pGSN may prime monocytes for antifungal responses while preventing excessive oxidative damage.

These results suggest that pGSN promotes a redox environment that is beneficial to pathogen clearance while minimizing collateral inflammatory stress, which may contribute to both enhanced monocyte function and reduced tissue injury observed *in vivo*.

## 4. DISCUSSION

Our study provides evidence that pGSN acts as a dual-function therapeutic agent in systemic *C. albicans* infection, both decreasing harmful inflammation and enhancing innate immune clearance. Using a murine model of candidemia combined with human monocyte assays, we demonstrate that exogenous pGSN reduces organ-specific pathology, lowers systemic cytokine expression, and improves the antifungal capacity of innate immune cells.

The organ-specific protection conferred by pGSN aligns with the known tropism of *C. albicans* during hematogenous dissemination. The kidneys are well-established as the primary target of fungal burden in murine models of candidemia, as well as in clinical settings (7). We observed widespread formation of fungal-rich microabscesses and granulocytic infiltration confined to the renal parenchyma. *C. albicans* is rarely found to be an etiological factor for pneumonia in clinical settings (26). In our study, lung pathology, although not associated with visible fungal cells, was characterized by emphysema, atelectasis, and the presence of hemosiderin-rich macrophages. These findings suggest that pulmonary damage may not be driven by direct fungal invasion but rather by a secondary inflammatory cascade (27). We have shown that pGSN treatment increases NO levels in monocytes while reducing ROS and H_2_S, suggesting a fine-tuned redox modulation that may be protective in inflammatory pulmonary environments. This interpretation is consistent with previously shown pGSN-driven NOS3 activation in inflamed lungs, which contributes to the resolution of inflammation (28). While NO production is essential for antifungal defense, its overproduction may contribute to tissue injury (29, 30). The context- and dose-dependent effects of NO underscore the need to balance reactive radical signaling for optimal host protection. In contrast, the liver remained largely unaffected in the acute 24-hour model, likely due to its robust populations of resident macrophages that efficiently clear pathogens from circulation (3). These observations underscore the significance of tissue-specific immune architecture in determining infection outcomes and therapeutic efficacy.

We administered pGSN subcutaneously, a route that offers practical advantages over intravenous delivery, especially in clinical settings. Subcutaneous injection enables sustained release and absorption, potentially prolonging the bioavailability of therapeutic proteins, such as pGSN (31). This delivery method is minimally invasive, easier to administer outside hospital settings. It has already been successfully tested in clinical trials involving critically ill patients (e.g., COVID-19), supporting its feasibility for broader therapeutic use (32).

While antifungal agents such as echinocandins and azoles directly target the fungal cell wall or membrane, they do not address the excessive inflammatory response that contributes to sepsis-related organ failure (33). Furthermore, antifungal resistance poses a significant threat, limiting therapeutic options (34). Our findings support the idea that pGSN, which has been evaluated for safety and efficacy in patients with infectious diseases (19, 21) could be deployed as an adjunctive, host-directed therapy in combination with standard-of-care antifungals.

In agreement with earlier studies reporting a rapid increase in TNF-α, IL-1β, IL-6, and chemokine gene expression (35) in human monocytes exposed to *C. albicans*, we observed a pronounced upregulation of pro-inflammatory genes, IL-1β, IL-6, TNF-α, and TGF-β, in the blood of mice infected with *C. albicans*, all of which were significantly inhibited by pGSN treatment. Notably, neither study observed a significant modulation of IFN-γ. However, we did detect TGF-β upregulation in infected mice, suggesting *that in vivo* signals are being generated from other cells.

We observed that pGSN significantly enhanced the phagocytic activity of human monocytes against *C. albicans*, as quantified by both the percentage of engaged cells and the phagocytic index. These findings build on previous work reporting a pGSN-driven boost in neutrophil-mediated killing of *Candida auris* (12). Moreover, our redox data expand upon observations showing that pGSN rebalancing of oxidative stress improves immune competence in human monocytes.

## 5. CONCLUSION

Our work has translational relevance for human candidemia, particularly in immunocompromised patients with impaired monocyte or neutrophil function. Limitations of this study include the use of immunodeficient mice, which precludes a full assessment of adaptive immunity. Additionally, the 24-hour endpoint, focused on innate immunity, restricts our understanding of long-term fungal clearance and tissue recovery. Future studies should explore the kinetics of pGSN effects over time, its combined effects with antifungal drugs, and its impact on other immunocompetent models or clinical settings.

In summary, our data position pGSN as a promising immunotherapeutic candidate in the management of fungal sepsis. Its dual ability to limit inflammation and enhance innate immune clearance provides a rationale for further preclinical and clinical investigation.

## ACKNOWLEDGMENTS

The authors are grateful to the staff of the Center for Experimental Medicine in Bialystok, Poland, for their technical assistance during the animal studies. This work was supported by the National Science Center, Poland, under Preludium bis 1 grant UMO-2019/35/O/NZ6/02807 (to R.B.) and Medical University of Białystok (B.SUB.25.403). Ł.S. was supported by the Foundation for Polish Science (FNP START 075/2024), by the Polish National Agency for Academic Exchange and Medical Research Agency (Walczak NAWA No. BPN/WAL/2023/1/00003/U/00001).

