## Supplemental materials for "Plasma Gelsolin Prevents Organ-Specific Inflammation and Enhances Innate Immune Function in a Systemic *Candida albicans* Infection"

**Table S1.** Primer pair sequence used in the study.

| Gene | Sequence (5′–3′) | |
|---|---|---|
| Il-1β | Forward | CAACCAACAAGTGATATTCTCCATG |
| | Reversed | GATCCACACTCTCCAGCTGCA |
| Il-6 | Forward | GAGGATACCACTCCCAACAGACC |
| | Reversed | AAGTGCATCATCGTTGTTCATACA |
| TNF-α | Forward | CATCTTCTCAAAATTCGAGTGACAA |
| | Reversed | TGGGAGTAGACAAGGTACAACCC |
| TGF-$\beta_1$ | Forward | TGACGTCACTGGAGTTGTACGG |
| | Reversed | GGTTCATGTCATGGATGGTGC |
| IL-10 | Forward | GGTTGCCAAGCCTTATCGGA |
| | Reversed | ACCTGCTCCACTGCCTTGCT |
| IFN-γ | Forward | TCAAGTGGCATAGATGTGGAAGAA |
| | Reversed | TGGCTCTGCAGGATTTTCATG |
| GAPDH | Forward | TGACCTCAACTACATGGTCTACA |
| | Reversed | CTTCCCATTCTCGGCCTTG |

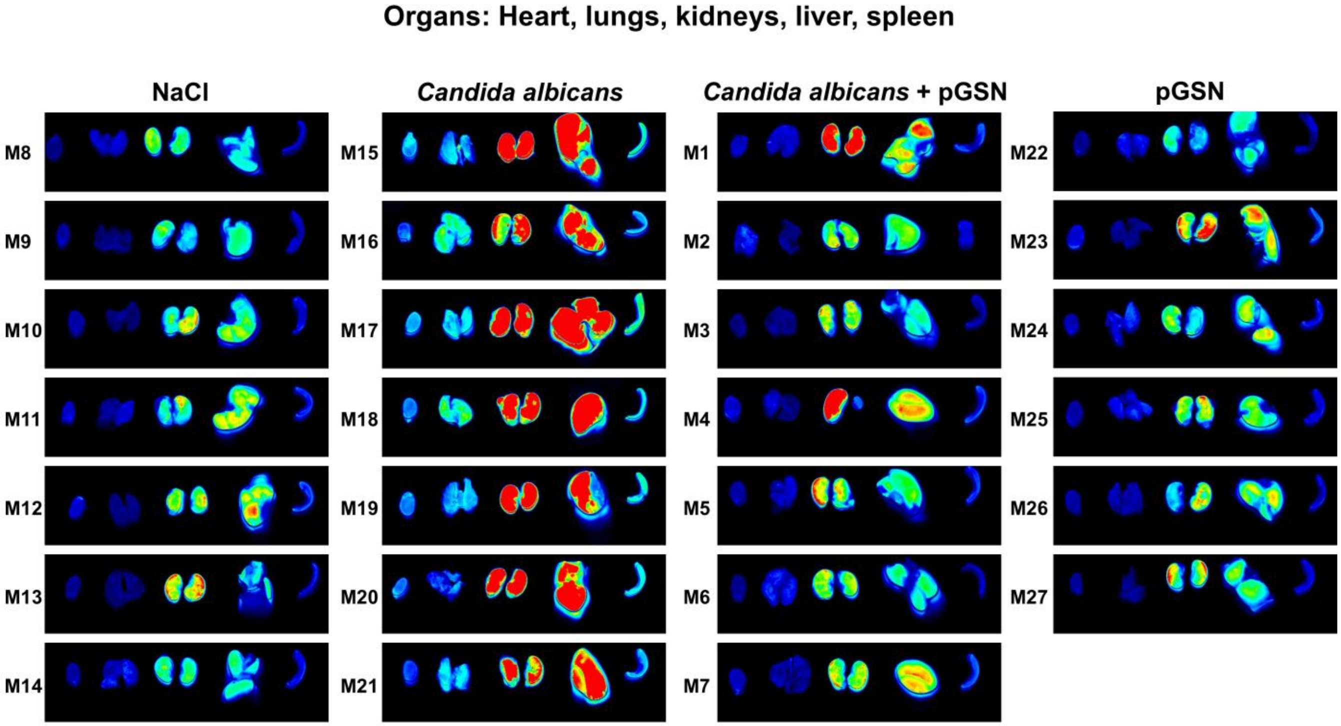


**Figure S1.** *Ex vivo* near-infrared fluorescence imaging of isolated organs (from the left: heart, lungs, kidneys, liver, and spleen) collected at 24 hours post-intravenous injection of *Candida albicans* or NaCl from all individual animals across the four experimental groups.

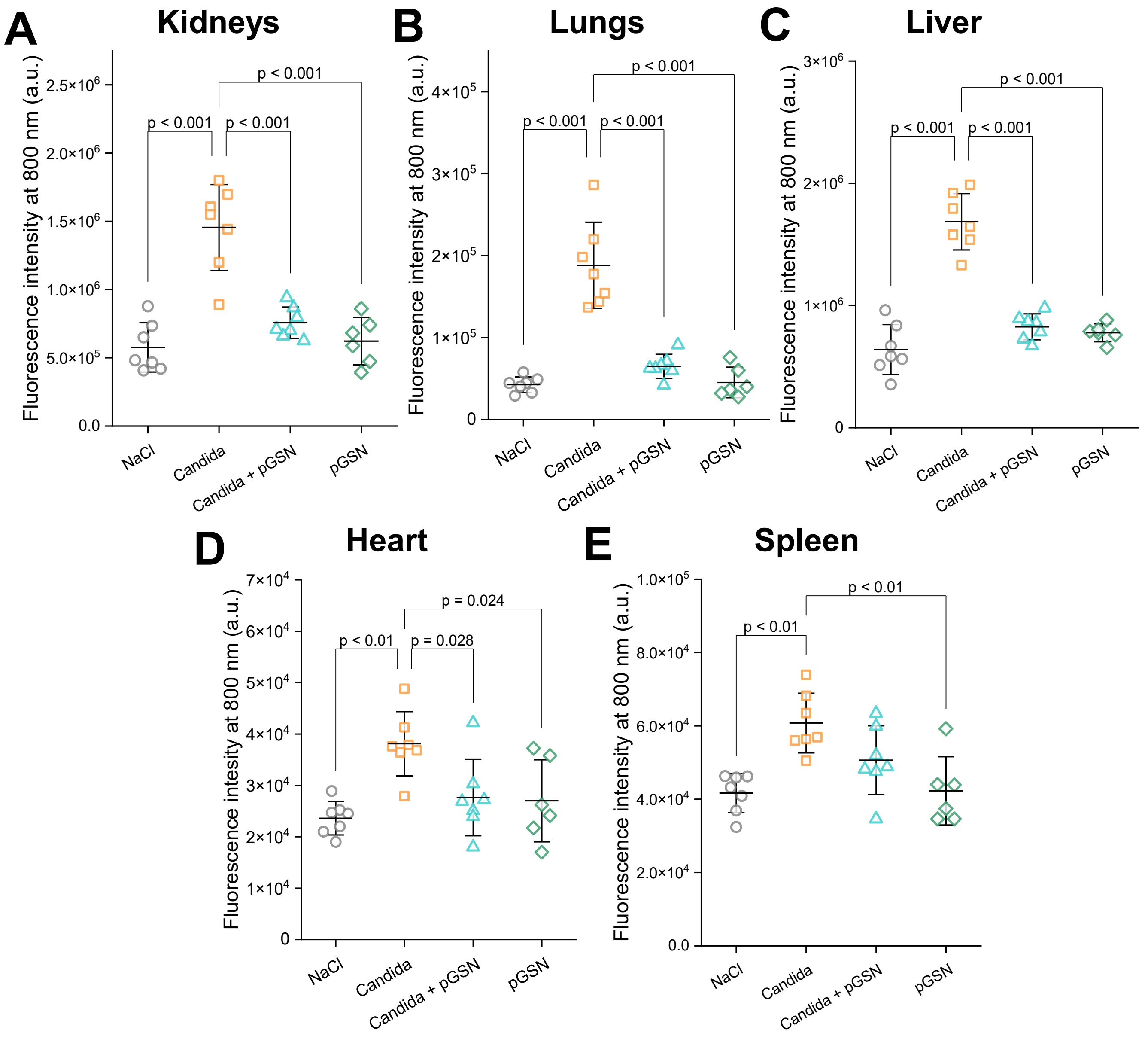


**Figure S2.** Individual animal-level organ fluorescence intensity data corresponding to Figure 2.*Ex vivo* near-infrared fluorescence intensities of IRDye® 800CW 2-deoxyglucose in isolated organs, kidneys (A), lungs (B), liver (C), heart (D), and spleen (E) from all individual animals across the four experimental groups. Each dot represents one animal. *Candida albicans* infection, which is attenuated by plasma gelsolin (pGSN) treatment. Data are presented as mean ± SD. Significance was assessed using one-way ANOVA with Tukey's post hoc test.

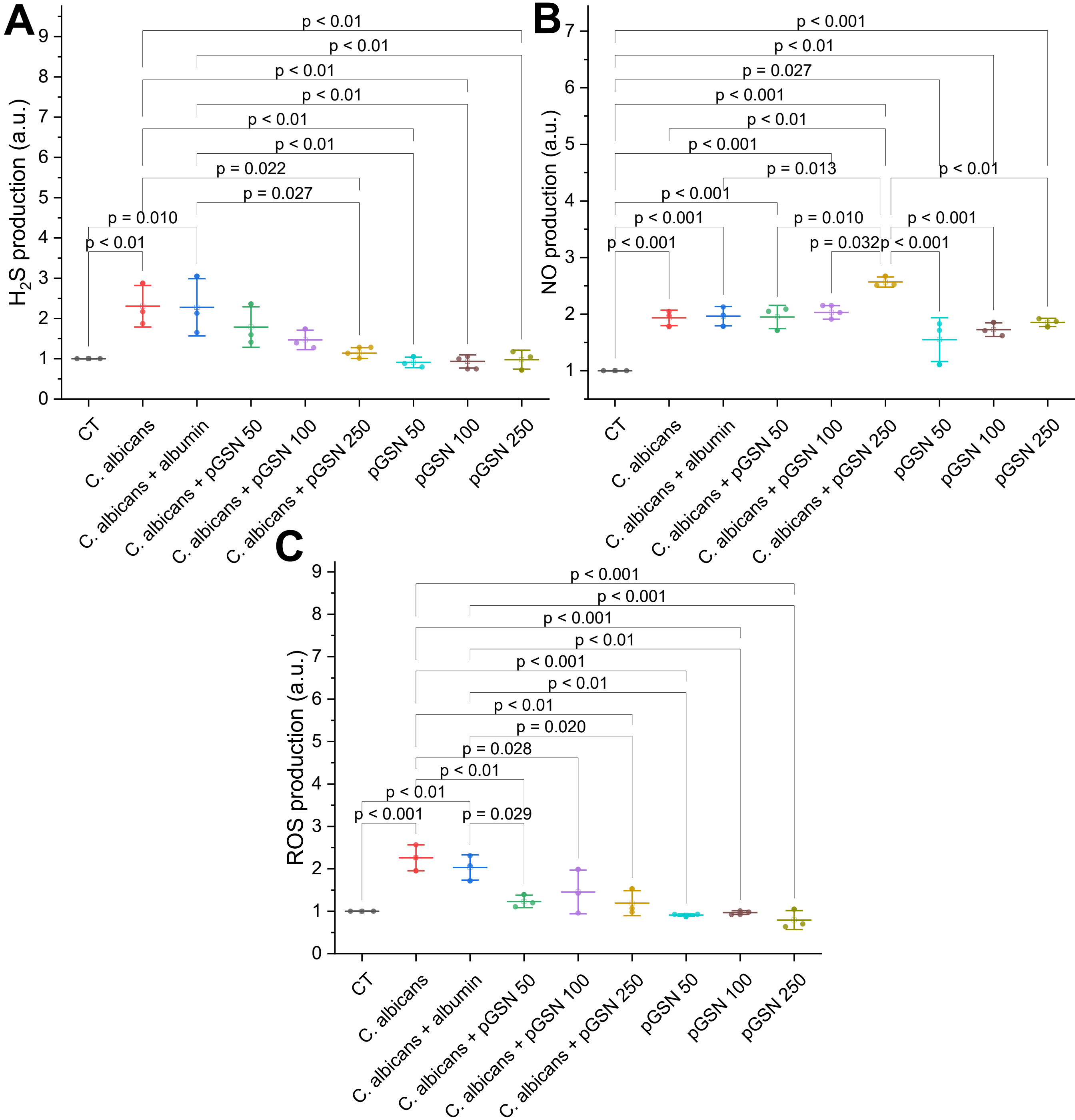


**Figure S3.** Plasma gelsolin modulates redox signaling in human monocytes in both basal and infected conditions. Monocytes from healthy donors were treated with Candida albicans, plasma gelsolin (pGSN), or both.(A) Hydrogen sulfide ($H_2S$),(B) nitric oxide (NO), and(C) reactive oxygen species (ROS) production were assessed using fluorescence-based assays. pGSN alone modestly enhanced NO levels without infection. Data are presented as mean ± SD from n =3. Statistical significance was determined by one-way ANOVA with Tukey's post hoc test.
